# Mapping the neural circuitry of cognitive restructuring in depressive and anxiety disorders

**DOI:** 10.64898/2026.07.12.738091

**Authors:** Alec J Jamieson, Trevor Steward, Kim L Felmingham, Christopher G Davey, Sevil Ince, James A Agathos, Bradford A Moffat, Rebecca K Glarin, Ben J Harrison

**Affiliations:** Department of Psychiatry, The University of Melbourne, Parkville, Australia; Melbourne School of Psychological Sciences, The University of Melbourne, Parkville, Australia; Centre for Mental Health and Brain Sciences, Swinburne University of Technology, Hawthorn, Australia; The Melbourne Brain Centre Imaging Unit, Department of Radiology, The University of Melbourne, Parkville, Australia

## Abstract

Cognitive restructuring, the process of identifying and challenging negative thoughts, is a key technique in the treatment of depressive and anxiety disorders. However, findings regarding its underlying neural circuitry remain inconsistent, potentially reflecting opposing effects of depressive and anxiety symptomatology that obscure disorder-specific dysfunction. A transdiagnostic sample of 73 clinical participants, predominantly with depressive or anxiety disorders, and 70 healthy controls completed a cognitive restructuring paradigm during 7 Tesla functional magnetic resonance imaging (fMRI). We undertook a multimodal characterisation of the restructuring process, integrating behavioural, physiological, and neuroimaging measures. Group-level fMRI analyses examined how depressive and anxiety symptom severity related to brain activation, while dynamic causal modelling characterised the directional neural influences between implicated regions. During challenging compared to repeating statements, greater depressive symptoms were associated with reduced left dorsolateral prefrontal cortex (dlPFC) activation. Conversely, greater anxiety symptoms were associated with greater dlPFC activation. Effective connectivity results revealed that depressive symptoms were associated with greater inhibition from the left ventrolateral prefrontal cortex (vlPFC) to the ventromedial prefrontal cortex, whereas anxiety symptoms were associated with greater excitation from the left dlPFC to amygdala and greater inhibition from the left vlPFC to amygdala. These symptoms additionally demonstrated a differential relationship with heart rate. While clinical participants modified negative beliefs as effectively as healthy controls, depressive and anxiety symptoms were associated with differential neural signatures during restructuring. This suggests that cognitive behavioural therapy may engage partially distinct mechanisms depending on symptom profile.

---

Cognitive theories of depression and anxiety posit that deeply ingrained negative beliefs are core features in the development and maintenance of these disorders (1). Once activated by subsequent stressors, these beliefs act to bias the processing of information (2-4). As a key component of cognitive behavioural therapy (CBT), cognitive restructuring represents a process that assists patients in identifying, evaluating, and modifying maladaptive automatic thought patterns (5). Through this, cognitive restructuring represents a key therapeutic mechanism underlying symptom improvement in depression and anxiety disorders. Given that altered self-beliefs are both a transdiagnostic feature and a key treatment target (6), identifying the neural mechanisms that enable their modification may provide critical insight into how psychological interventions produce therapeutic change.

Growing evidence indicates that cognitive restructuring involves complex interactions between brain systems implicated in self-belief processing and affect regulation (7, 8). The ventromedial prefrontal cortex (vmPFC), as a core hub of the default mode network, is central to these functions, including supporting cognitive representations of the self (9). This role in self-referential processing is thought to reflect the vmPFC’s broader function in encoding the subjective and affective value of stimuli (10-12). Affect regulation also appears to be important for successful restructuring, as heightened negative affect limits the integration of novel positive information needed to revise negative beliefs (13, 14), thereby constraining belief updating. This regulation is supported by the cognitive control network, which contributes to the metacognitive process that allows self-representations to become the objects of attention (15). Within this network, the presupplementary motor area (preSMA) is hypothesised to aid in strategy selection by modulating lateral prefrontal regions through its roles in directing internal attention (16) and inhibitory control (17). The dorsolateral prefrontal cortex (dlPFC) is subsequently suggested to maintain and manipulate strategy-relevant representations and regulatory goals in working memory (18), while the ventrolateral prefrontal cortex (vlPFC) is more directly implicated in suppressing affective responses during reappraisal (19). The regulatory influence of lateral prefrontal regions is, in turn, thought to act on the amygdala (20). The relatively sparse anatomical connectivity between the lateral prefrontal cortex and the amygdala (21-23), however, indicates that intermediary regions including the vmPFC also play an important role in mediating this regulation (24-26).

Given the central role of cognitive restructuring in the treatment of anxiety and depressive disorders, growing research has sought to characterise dysfunction within this circuitry and its relationship with symptom expression. Accordingly, widespread prefrontal alterations have been shown across depressive and anxiety disorder studies investigating representations of the self (6). Specifically, depression appears to be more strongly associated with increased anterior cingulate cortex activation, whereas anxiety is associated with increased insula and dlPFC activation (6). The clinical relevance of these activation patterns is highlighted by the fact that CBT has been shown to normalise hyperactivation across the dorsal anterior cingulate, dlPFC, and insula in anxiety disorders (27). Conversely, the cognitive reappraisal and emotion regulation literature has reported reduced activity in the dlPFC, vlPFC, and dorsal anterior cingulate cortex across both anxiety and depressive disorders (28, 29), although more recent work has also yielded inconsistent findings (30, 31). These inconsistencies may reflect the opposing effects of depressive and anxiety symptoms being masked at the disorder level due to the high rates of comorbidity between these conditions (32-34). Such symptom divergence has previously been observed during interoceptive processing, where anxiety and depressive symptoms demonstrated opposing neural associations (35), highlighting the need for dimensional approaches to resolve apparent neural heterogeneity.

To address this, we examined the neurocircuitry of cognitive restructuring in a transdiagnostic clinical sample, predominantly with anxiety and depressive disorders, using 7-Tesla (7T) fMRI and a novel cognitive restructuring task (36). We undertook a multimodal characterisation of the task, integrating behavioural, physiological, and neuroimaging measures to provide a comprehensive account of the restructuring process. We assessed task-evoked activation and used dynamic causal modelling (DCM; 37) to characterise how depressive and anxiety symptoms were differentially associated with regional activity and directional influences among prefrontal-subcortical regions. We hypothesised that depressive symptoms would be associated with hypoactivation of the dlPFC and vlPFC (28, 29) and reduced inhibitory connectivity from lateral prefrontal regions to the amygdala (38), with the vmPFC serving as the principal mediating node. Conversely, we hypothesised that anxiety symptoms would be associated with greater activity in the insula, amygdala, and dlPFC (6), alongside greater inhibitory prefrontal-to-amygdala connectivity (39), also routed through the vmPFC.

## Method

### Participants

A transdiagnostic sample of 82 clinical participants with symptoms of depression and anxiety was recruited from The University of Melbourne Psychology Clinic and the School of Psychological Sciences, as described in a previous publication (40). Participants were assessed using the Diagnostic Interview for Anxiety, Mood, and OCD and Related Neuropsychiatric Disorders (41) to identify those who met DSM-5 diagnostic criteria for a depressive and/or anxiety disorder. Participants meeting criteria for a stress and/or eating disorder were also included, given the high level of depressive and anxiety symptoms present in these conditions (42, 43). Exclusion criteria for this group included meeting criteria for psychosis, bipolar disorder, paraphilic disorder, dissociative disorder, conduct disorder, or being actively suicidal.

In addition, we recruited 88 healthy control participants through online advertisements. These participants had no current or past diagnosis of a mental health disorder as assessed through the Mini-International Neuropsychiatric Interview-7 (44). All participants included in this study were aged 18 to 40 years, fluent English speakers, had no MRI contraindications, and possessed normal or corrected-to-normal vision. Prior to participation, and following a complete description of the study protocol, all participants provided written informed consent. The University of Melbourne Human Research and Ethics Committee approved this study and its procedures. To assess symptom severity, the Depression, Anxiety, and Stress Scale (DASS) was administered, and subscale scores for depression, anxiety, and stress-tension were obtained (45).

Of the original sample, twenty-seven participants were excluded: eight for technical issues during acquisition (three controls, five clinical), ten for non-completion or incorrect completion of the fMRI paradigm (eight controls, two clinical), and nine for excessive head movement during scanning (seven controls, two clinical; see Supplementary Methods for details). The final sample comprised 70 healthy controls and 73 clinical participants.

### Cognitive restructuring paradigm

Participants completed a previously detailed cognitive restructuring paradigm (8, 36), which leveraged Socratic questioning to challenge negative self-perceptions, akin to skills employed in cognitive psychotherapy (46). Prior to scanning, participants were trained in cognitive restructuring techniques including active rebuttal, reinterpretation, and perspective shifting. Once participants’ proficiency was determined by the research staff, the paradigm was administered during concurrent fMRI scanning. The cognitive restructuring task consisted of a single run consisting of 16 blocks (*Supplementary Fig. S1*). Each block began with the on-screen presentation of a negative self-cognition statement drawn from CBT and psychopathology literature (Supplementary Table S4)(47, 48). Participants then indicated via button press whether they would repeat or restructure the given statement using their pre-learned strategies, with instructions to balance the two across the run. After 9 seconds, the statement reappeared and participants engaged in their chosen strategy for 12 seconds: either restructuring the statement (Challenge condition) or repeating it (Repeat condition). This was followed by a fixation period, which occurred for an average of 6 seconds, after which the next block began with a new statement. The paradigm was developed and presented using E-Prime 3.0 software (Psychology Software Tools, Pittsburgh, PA).

Before and after the task, all participants rated their endorsement of each of the 16 negative statements on a 7-point self-report scale (1 = strongly disagree; 7 = strongly agree). To assess the effectiveness of cognitive restructuring, pre- and post-scan endorsement levels of negative self-beliefs were compared between groups using a mixed-model analysis of variance (ANOVA), with time (pre vs post) as the within-subjects factor and group as the between-subjects factor. We additionally investigated heart rate during the task to assess the influence of task conditions on the functioning of the autonomic nervous system (see Supplementary Materials for details). Heart rate is a sensitive marker of autonomic arousal during emotional and cognitive processing (49). Altered physiological responses, including heart rate, to emotional and psychological stressors have also been observed in individuals with elevated depressive symptoms (50). A linear mixed-effects model was therefore used to examine the effects of condition and depressive and anxiety symptom severity on heart rate.

### General linear modelling

Following preprocessing of the fMRI data (see Supplementary Methods), first-level general linear model (GLM) analyses were conducted for each participant using SPM12. Regressors for the challenge, repeat, and decide conditions were specified using the onsets and durations of each block, convolved with a canonical haemodynamic response function. Physiological noise regressors were then added to the GLM, and a high-pass filter (1/128 seconds) was applied to account for low-frequency drifts. The FAST method was used to estimate temporal autocorrelations due to its enhanced performance at short repetition times (51). Primary contrast images from each participant (Challenge > Repeat and Challenge < Repeat) were carried forward to second-level analyses using the summary statistics approach to random-effects analyses. At the second level, we modelled the depressive and anxiety DASS subscales as separate regressors within a single GLM to isolate the effects of each symptom domain, adjusting for age, sex, and antidepressant medication use as covariates of no interest. To ensure that collinearity between regressors did not bias the estimates, we examined the variance inflation factor (VIF) of the depression and anxiety DASS subscales. This revealed a VIF of 1.76 between the depression and anxiety subscales. As this value is under 5, this indicates an acceptable level of collinearity between factors for modelling (52).

### Dynamic causal modelling

DCM estimates the rate of change in the activity of a region as a function of activity in another (i.e., effective connectivity, measured in hertz; Hz). In task-based DCM, connectivity is decomposed into a baseline component (intrinsic connectivity) and a component modulated by the experimental condition of interest (53). Positive values are indicative of a putative excitatory influence between regions, while negative values represent a putative inhibitory influence. Self or within-region connections are conversely represented by unitless log scaling parameters that modify the default inhibitory connectivity of -0.5 Hz. For these self-connections, positive values for these parameters are indicative of greater inhibition, whereas negative values represent reduced inhibition.

### Timeseries extraction, model specification, estimation, and inference

Our model space for DCM consisted of five regions of interest: the preSMA, dlPFC, vlPFC, vmPFC, and amygdala (*Supplementary Fig. S2*). These regions were chosen based on prior evidence implicating them in cognitive restructuring for healthy individuals (7, 36). Representative timeseries were extracted from each region at the subject-level following published guidelines (53). For full details of the extraction methodology, see Supplementary Methods. The candidate model was specified using DCM version 12.5 in SPM12. Intrinsic connectivity was modelled as bidirectional between all regions, except between the preSMA and amygdala, which was assumed to be mediated via other prefrontal areas, given their lack of anatomical connection. Modulation by the Challenge condition was also specified across these connections. The ‘Task’ condition (onset of all Challenge and Repeat blocks) was defined as the driving input into the preSMA, given this region drove the largest input effect in prior investigations (36). This input was additionally mean-centred, so that the intrinsic connectivity represented the average connectivity. Parametric empirical Bayes (PEB) was then used to estimate symptom-related effects on effective connectivity (for additional estimation and inference details see Supplementary Methods). Leave-one-out cross-validation (LOOCV) in the PEB framework was finally used to evaluate the ability of connectivity parameters to predict individual differences in symptoms (Supplementary Methods and Results).

## Results

### Demographic and behavioural results

No significant between-group differences were observed in age, sex, or handedness (Table 1). As expected, clinical participants exhibited significantly higher symptom severity than healthy controls across all three DASS subscales. Mann–Whitney U tests indicated significant group differences for depression scores (*U* = 4807.50, *z* = 9.12, *p* < .001), anxiety scores (*U* = 4566.50, *z* = 8.17, *p* < .001), and stress scores (*U* = 4586.00, *z* = 8.22, *p* < .001).

**Table 1.**
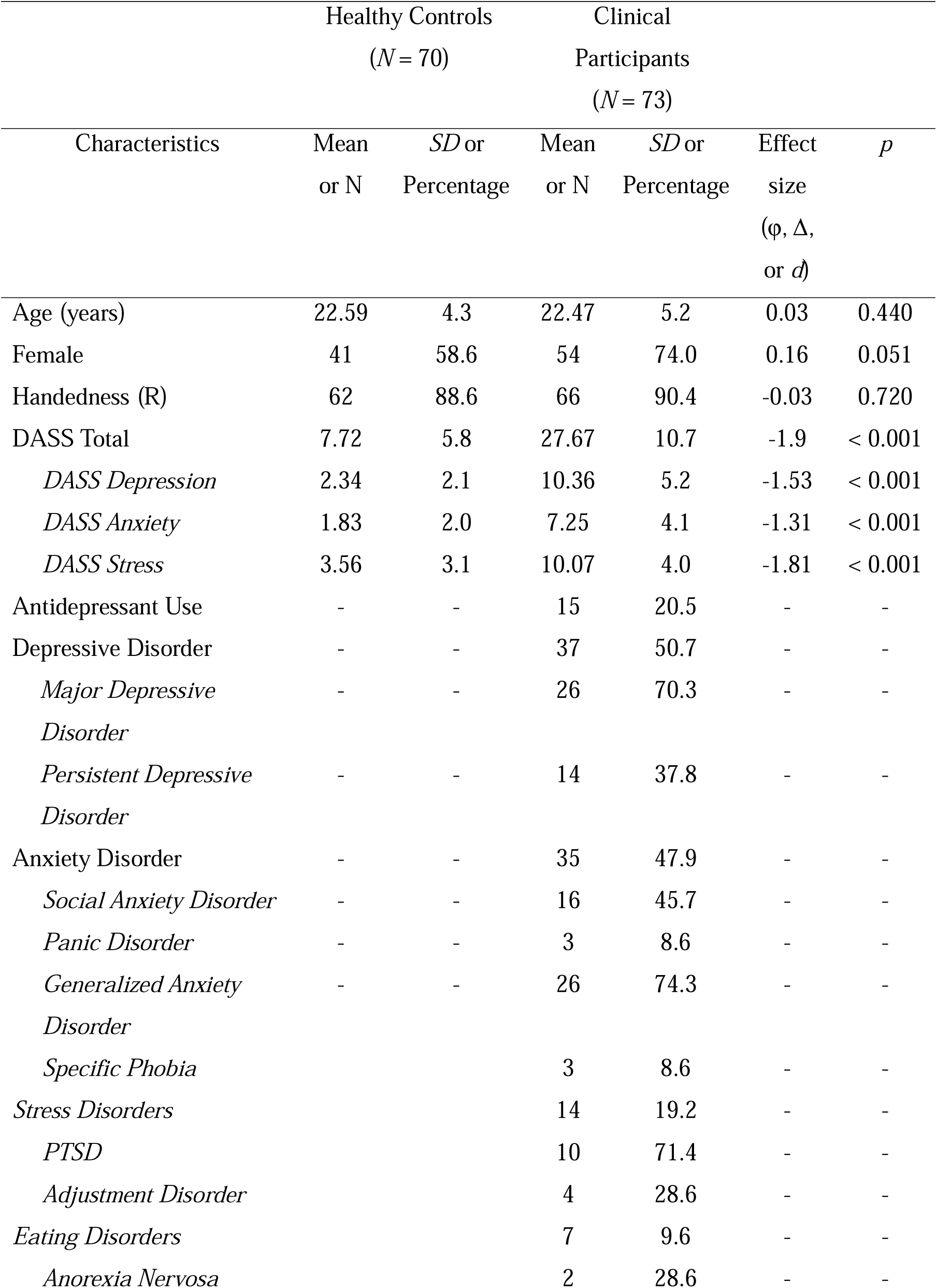
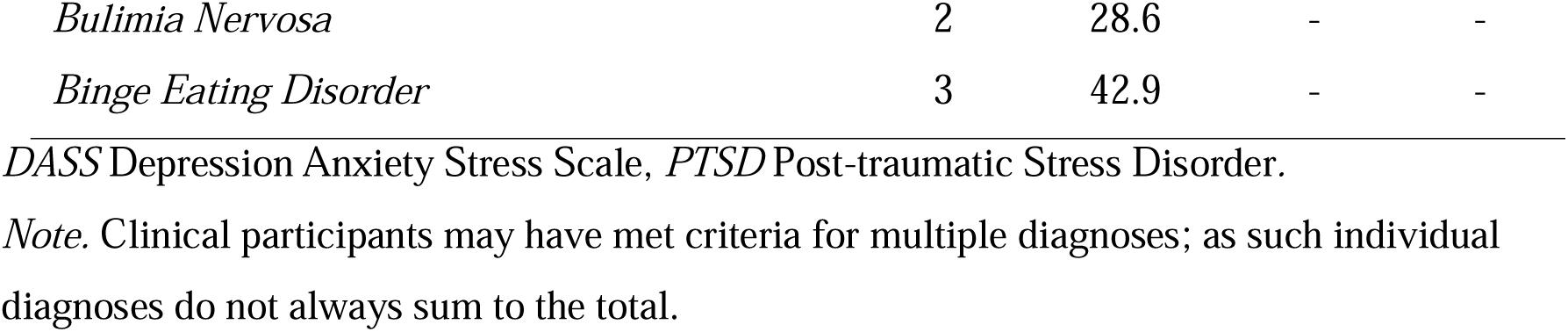
Comparison of Characteristics Between Healthy Controls and Clinical Participants.

Before scanning, participants rated their endorsement of each negative statement used in the task. Clinical participants endorsed most statements more strongly than healthy controls (Supplementary Results; *Supplementary Fig. S3*). After scanning, participants re-rated their endorsement of the negative statements. Endorsement of challenged statements decreased significantly from pre- to post-paradigm (*F*(1, 140) = 159.19, *p* < .001), indicating that cognitive restructuring was successful. Although clinical participants endorsed statements more strongly at both timepoints (*F*(1, 140) = 93.88, *p* < .001), the group × time interaction was not significant (*F*(1, 140) = 1.39, *p* = .240), indicating that the magnitude of restructuring-related reduction did not differ between groups (*Fig. 1*). There were no significant correlations between pre-to-post changes in endorsement and either depression or anxiety symptoms (*p* = .104 and *p* = .070, respectively).

**Fig. 1.**
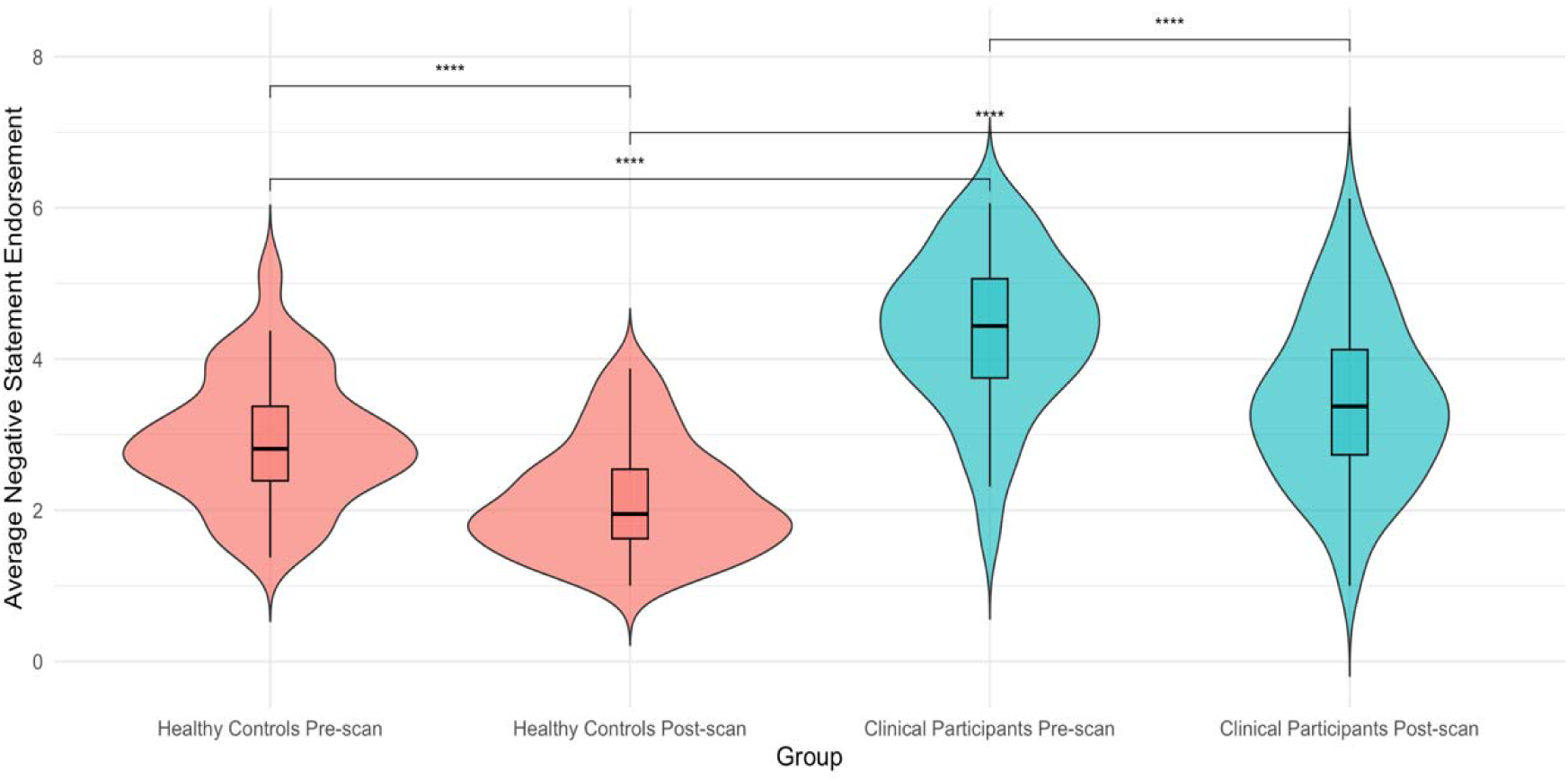
Violin plots depicting the change in challenged negative statement endorsement scores from pre-scan to post-scan. Results show a significant difference between healthy controls and clinical participants at both time points, as well as significant reductions following challenging. Stars represent statistical significance after Bonferroni correction for multiple comparisons, with **** representing *p* < .0001.

### General linear modelling results

We first characterised the broad activation pattern associated with cognitive restructuring by examining regions more active during challenging than repeating negative statements (Challenge > Repeat). Consistent with our previous work (8, 36, 54), we observed increased activity in the preSMA extending to the dorsal anterior cingulate cortex, bilateral caudate nucleus extending to the dorsomedial thalamus, cerebellum (right posterior lobe and vermis), left supplementary motor area, left posterior dlPFC extending to the vlPFC, posterior cingulate cortex extending to the retrosplenial cortex, bilateral temporal pole, vmPFC, and left inferior parietal lobule (*Fig. 2*).

**Fig. 2.**
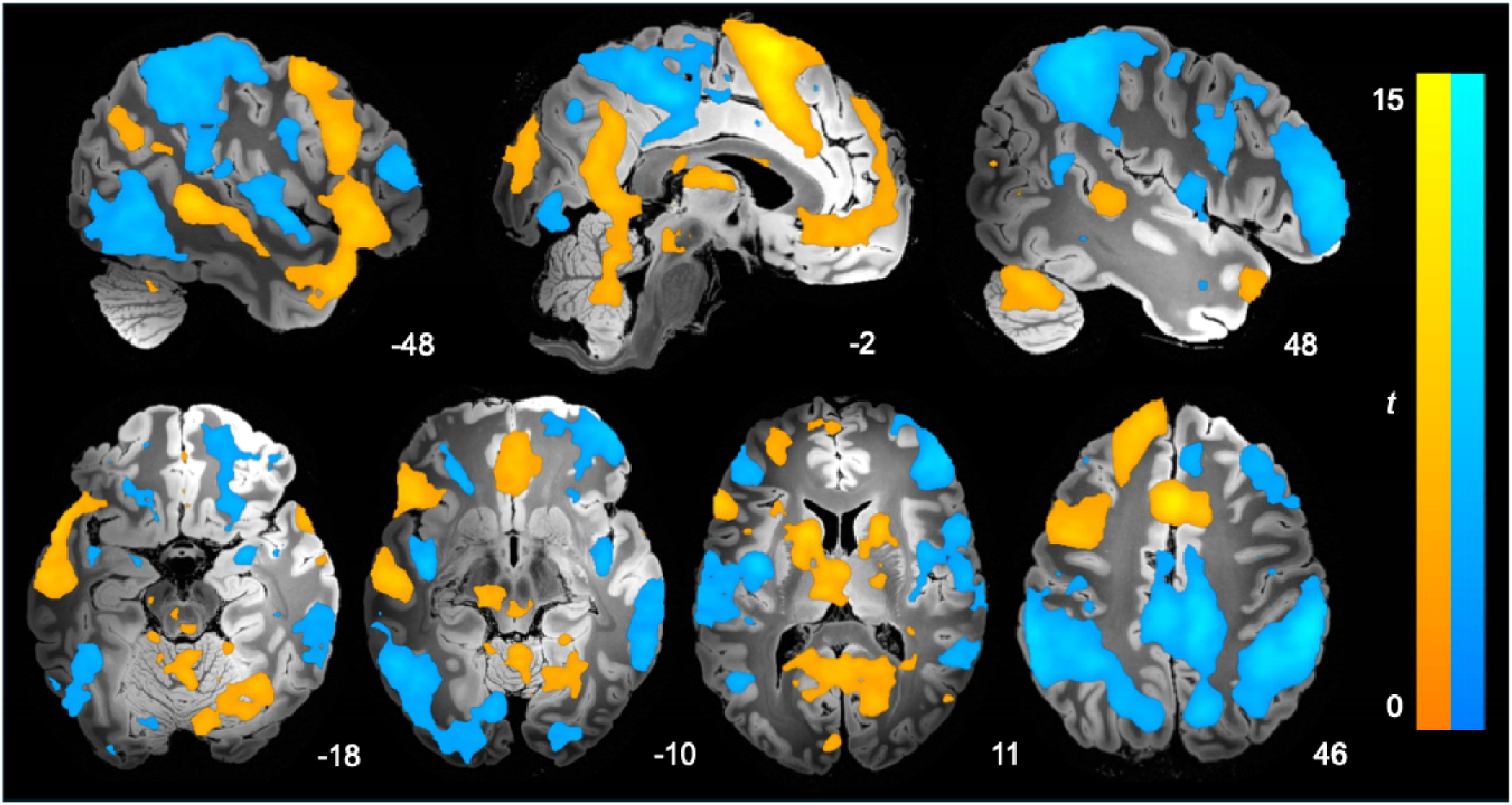
Significant activation during challenging compared with repeating negative statements. Shown are one-sample t-tests across all participants for Challenge > Repeat (warm) and Challenge < Repeat (cool). Results were thresholded at *p*_FDR_ < .001. Colour bars represent t-statistics. Results visualised on the “Synthesized_FLASH25” (500 μm, MNI space) ex vivo template (87). Left = Left.

Conversely, several brain regions displayed greater deactivation during challenging compared to repeating negative statements (or alternatively stated, were more active during repeating than challenging; Challenge < Repeat). These regions included the bilateral angular gyrus extending to the posterior operculum, dorsal posterior cingulate cortex/precuneus, left fusiform gyrus, right anterior vlPFC, bilateral posterior insula, and the amygdala (*Fig. 2*). For full results, see Supplementary Tables S1 and S2.

### Associations with symptom severity

Within the same model, we examined how the depressive and anxiety DASS subscales were associated with brain activation during cognitive restructuring (*Fig. 3*). Depressive symptoms were *negatively* associated with activation of the left dlPFC (peak: x=-48, y=22, z=27; cluster size = 157; *t* = 4.52). In overlapping voxels of the left dlPFC, anxiety symptoms showed the *opposite* pattern, being *positively* associated with activation (peak: x=-45, y=21, z=19; cluster size = 72; *t* = 4.05; all clusters *P_FDR_* < .05, cluster-forming threshold *p* < .001; *Fig. 3*). The association between dlPFC activation and both depression and anxiety symptoms was also evident within the clinical group alone (*Supplementary Fig. S4*).

**Fig. 3.**
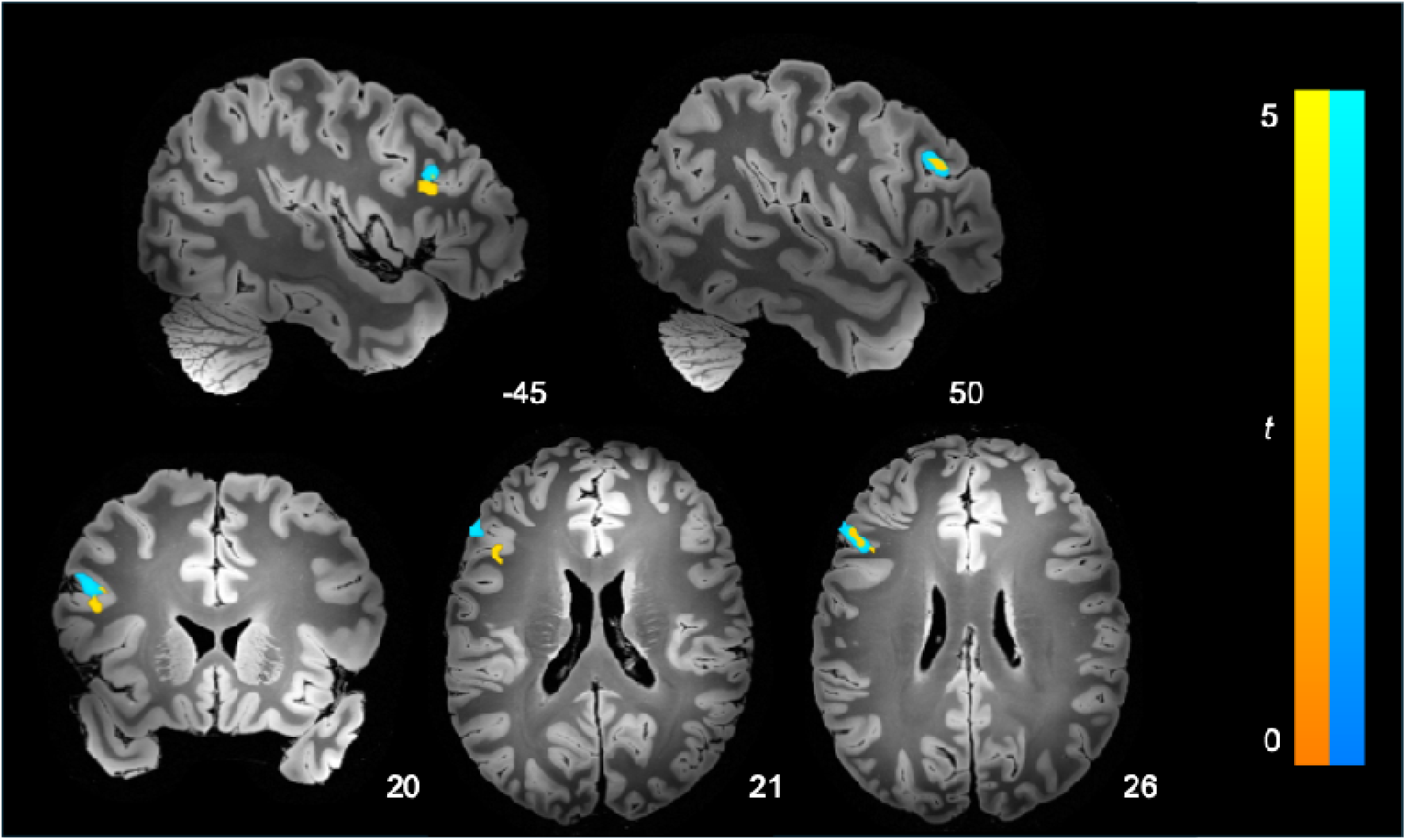
Significant associations with depressive and anxiety symptom severity and brain activation during challenging compared with repeating negative statements. Positive associations with anxiety symptom severity (warm) and negative associations with depressive symptom severity (cool). *P* < .001, FDR cluster-wise corrected. Colour bars represent t-statistics. Results visualised on the “Synthesized_FLASH25” (500 μm, MNI space) ex vivo template (87). Left = Left.

### Network modulation during cognitive restructuring

We next examined how these symptoms related to prefrontal–subcortical dynamics during cognitive restructuring. Connectivity was estimated both during the challenging of negative statements and intrinsically (for intrinsic results see Supplementary Materials and *Supplementary Fig. S5*).

Across all participants, challenging negative statements was associated with strong excitatory modulation from the preSMA to the dlPFC, vlPFC and vmPFC, and from the amygdala to dlPFC and vlPFC (*Fig. 4A*). Inhibitory modulation was observed from the dlPFC to vlPFC and vmPFC, from the amygdala to vmPFC, and from both the vlPFC and vmPFC to all other regions. Challenging negative statements, therefore, engaged both ‘top-down’ and ’bottom-up’ pathways linking the preSMA and amygdala, though the dominant pattern was one of prefrontal-driven inhibition of the amygdala.

**Fig. 4.**
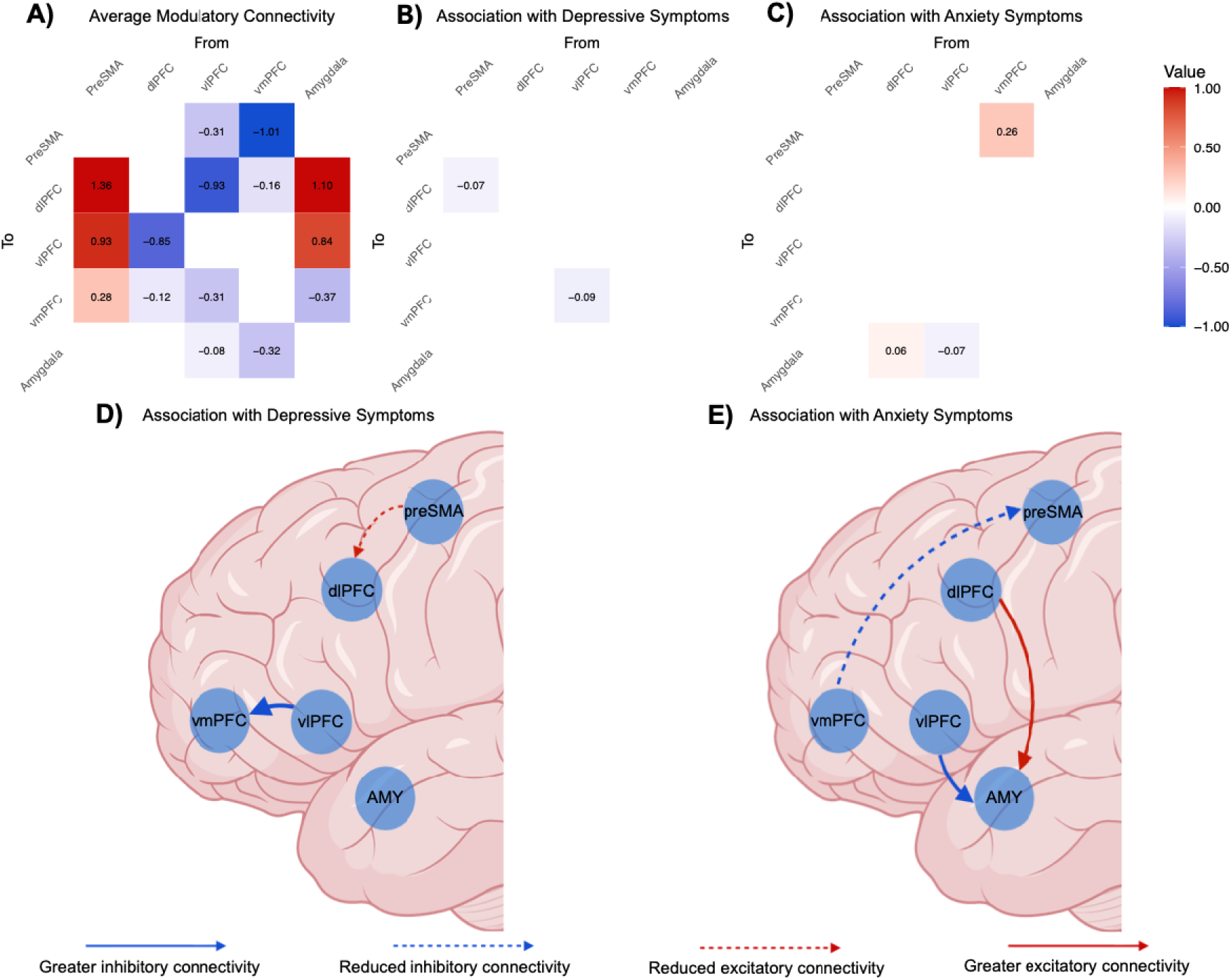
Challenge related modulation of effective connectivity across healthy controls and clinical participants. Adjacency matrices show the average effects observed in **A)** all participants, **B)** the effect of depressive symptoms, and **C)** the effect of anxiety symptoms. For **A)** red cells indicate excitatory connections, whereas blue cells indicate inhibitory connections. For **B)** and **C)** these values add (red) to or subtract (blue) from the connectivity in **A)**. Cells representing connectivity between two regions are measured in Hz and diagonal cells indicate inhibitory self-connections, which are unitless log scaling parameters. **D)** and **E)** illustrate effective connectivity parameters associated with depressive and anxiety symptoms, respectively. Image created with BioRender (www.biorender.com).

Depressive symptoms were associated with greater inhibition from the vlPFC to vmPFC and reduced excitation from the preSMA to dlPFC during challenging (*Fig. 4B* and *D*). Conversely, anxiety symptoms were associated with reduced inhibition from the vmPFC to the preSMA, greater excitation from the dlPFC to the amygdala, and greater inhibition from the vlPFC to the amygdala (*Fig. 4C* and *E*). In summary, depressive symptoms were primarily associated with altered connectivity between prefrontal regions, whereas anxiety symptoms were associated with a mixed pattern of reduced prefrontal inhibition and enhanced ’top-down’ cortical influence on the amygdala. For examination of these relationships in only the clinical group, see Supplementary Materials (*Supplementary Fig. S6*).

### Heart rate results

To examine whether these symptom profiles were also associated with differences in autonomic arousal, we examined heart-rate responses across the task. Heart rate showed a temporal and condition-related pattern, varying across the task and between the Repeat and Challenge conditions (*Fig. 5A–B*). Across the sample, there was a robust main effect of condition on heart rate, such that heart rate was significantly higher during the Repeat condition relative to Challenge (*b* = −1.18, *t* = −31.48, *p* < .001). There was also a significant main effect of time, indicating a gradual decrease in heart rate over the course of the trial (*b* = −0.07, *t* = −6.23, *p* < .001). Neither anxiety nor depression showed significant main effects on heart rate (*b* = −0.33, *t* = −0.24, *p* = .808; *b* = 0.54, *t* = 0.40, *p* = .694, respectively), however, significant interaction effects were observed for both symptoms (*Fig. 5C*). Specifically, higher depressive symptoms were associated with a greater condition effect on heart rate (*b* = −0.22, *t* = −4.38, *p* < .001), indicating greater heart rate differences between Challenge and Repeat as depression increased. In contrast, anxiety symptoms showed a significantly reduced condition effect (*b* = 0.18, *t* = 3.65, *p* < .001), suggesting that higher anxiety severity attenuated condition-related differences in heart rate. For full results, see Supplementary Table S3. These results were consistent when analysing just the clinical participants (*Fig. 5D*; Supplementary Materials).

**Fig. 5.**
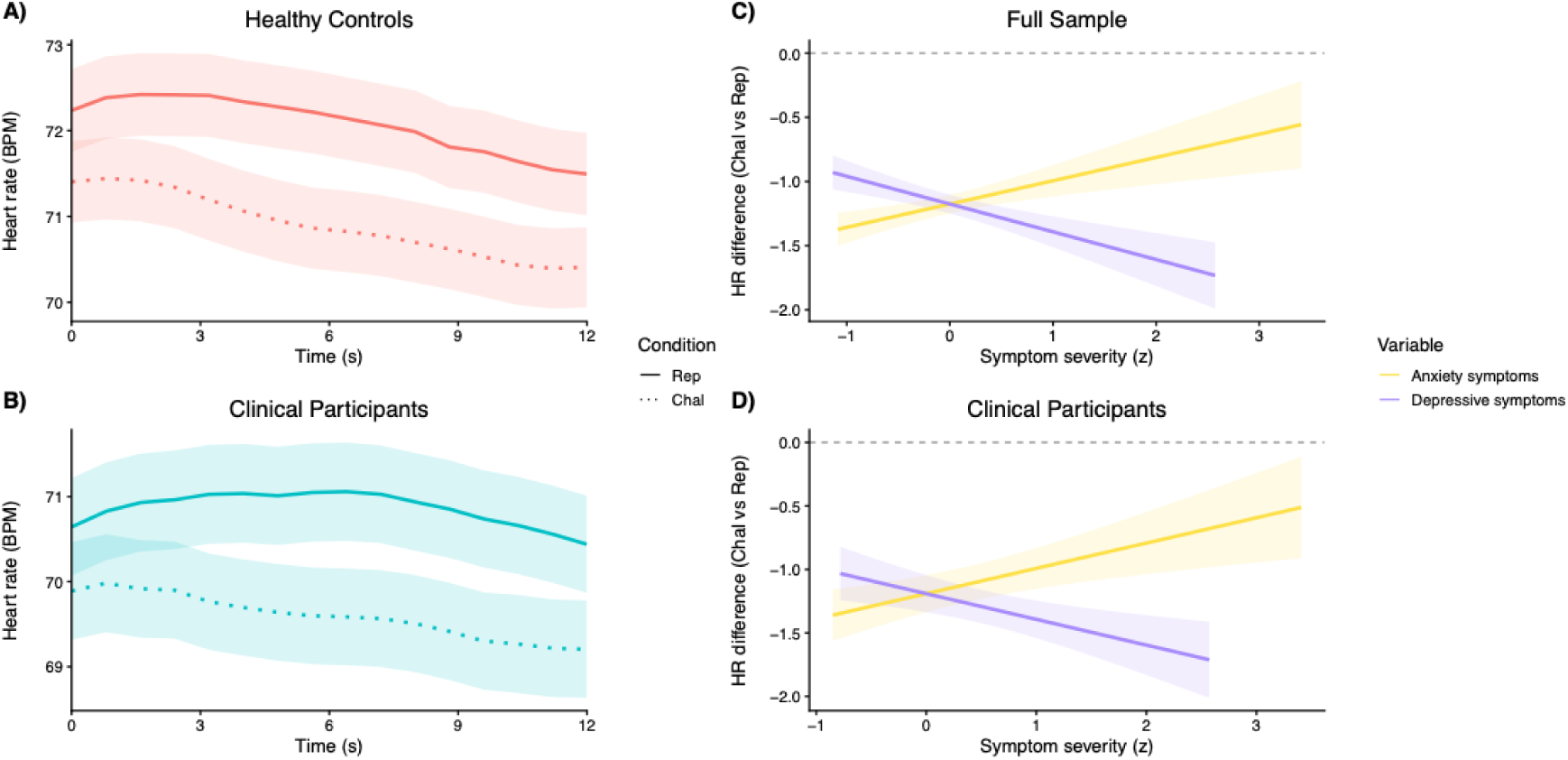
Task-related and symptom-modulated changes in heart rate. Mean heart rate over time during the Repeat (Rep) and Challenge (Chal), stratified by **A)** healthy controls and **B)** clinical participants, with shaded bands representing the standard error of the mean. Model-derived estimates of the differential effect of condition (Challenge vs Repeat) on heart rate as a function of symptom severity for **C)** the full sample and **D)** just the clinical participants. Lines represent predicted effects from the linear mixed-effects model. Positive values indicate a higher heart rate in the Challenge condition relative to the Repeat condition.

## Discussion

This study examined the neurocircuitry of cognitive restructuring and its links to depressive and anxiety symptoms. Challenging negative statements engaged the dlPFC differently by symptom type: anxiety was associated with increased activation, depression with reduced activation. Depressive symptoms were also linked to altered preSMA to dlPFC and vlPFC to vmPFC connectivity, while anxiety showed altered connectivity between prefrontal regions and the amygdala. Heart rate data also supported this divergence, with depressive symptoms increasing the difference between challenging and repeating, whereas anxiety symptoms reduced this difference. Together, these findings show that although clinical participants restructured negative statements as efficiently as healthy controls, depression and anxiety were marked by dissociable patterns of prefrontal-subcortical circuitry and autonomic regulation.

### Anxiety and depressive symptom associations with the posterior dlPFC

We observed that activation associated with depression and anxiety symptoms during restructuring was localised to the posterior portion of the dlPFC. The dlPFC is involved in monitoring and manipulation of working memory contents (55) and exhibits a posterior-to-anterior functional gradient (56). Within this gradient, the posterior dlPFC tracks dynamic changes in the environment and individual, and the anterior dlPFC supports more abstract representations (57, 58). The posterior dlPFC is therefore thought to support the integrative process underpinning cognitive control (59). Evidence indicates that posterior dlPFC activity reflects the cognitive effort required to overcome a heightened affective load and implement cognitive control, with reappraisal of high-intensity emotions eliciting greater activation than low-intensity emotions (60). Consistent with this role, transcranial magnetic stimulation (TMS) targeting peak dlPFC activation during cognitive restructuring accelerated heart rate regulation and increased subsequent use of cognitive restructuring, despite not reducing overall distress (61). Together, these findings implicate the posterior dlPFC as a key locus for the efficient instantiation of cognitive control, particularly while under affective load.

Expanding on this work, we observed that posterior dlPFC activation during restructuring tracked anxiety and depressive symptoms in opposite directions. Anxiety and depression primarily differ phenomenologically in their temporal focus (future versus past) and content specificity (threat versus hopelessness/helplessness)(62). For anxiety in particular, Attentional Control Theory proposes that threat-related stimuli capture attentional resources, increasing the cognitive effort required for regulating non-threat content (63, 64). Anxiety is therefore hypothesised to impair processing efficiency rather than performance effectiveness (64), with those with anxiety having preserved performance by investing compensatory effort (64), which manifests as increased dlPFC activation (65) and greater affect regulation.

In contrast, depressive symptoms were associated with hypoactivation of the dlPFC, a pattern previously interpreted as impaired inhibition of negative information (66, 67). Such inhibitory failures have been thought to reflect the intrusion of self-related thought into task-relevant processing, consistent with established links between inhibitory deficits and rumination (68). Notably, both reduced rumination and increased self-efficacy have been identified as mediators of CBT effectiveness for social anxiety (69) and depressive disorders (70, 71). Although the capacity for cognitive restructuring appears to be preserved across depressive and anxiety disorders (72), it is not enacted in relevant contexts (73). As such, these findings suggest that the primary constraint on cognitive restructuring in depressive and anxiety disorders is not the capacity for implementation, but the initiation and efficient deployment of the process in the face of competing demands. Similar dissociations between anxiety, depression, and dlPFC function have been reported during recall of negative events (74) and the explicit processing of negative emotional expressions (75), indicating that these alterations generalise across tasks requiring the manipulation of negative stimuli in working memory.

### Network associations with anxiety symptoms

Anxiety symptoms were also associated with enhanced influences from lateral prefrontal regions to the amygdala during restructuring negative statements, alongside altered intrinsic connectivity involving these same regions. Given the well-established link between anxiety and exaggerated threat appraisal (5, 76), there has been substantial neuroscientific focus on the amygdala (77). Despite this focus, associations between amygdala function and anxiety have been inconsistent (77). This inconsistency likely reflects multiple factors, including the amygdala’s complex functional architecture (77) as well as compensatory mechanisms that downregulate altered amygdala responses. Although our activation analyses did not reveal an association between amygdala activation and anxiety, the connectivity analyses revealed both greater intrinsic self-inhibition of the amygdala and greater modulatory influences from the vlPFC and dlPFC to the amygdala during restructuring. This pattern is consistent with a compensatory role for the lateral prefrontal regions, whereby their enhanced influence over the amygdala may support successful belief changes in those with greater anxiety symptoms (78). Although this finding is consistent with the proposed regulatory role of these regions, the lateral prefrontal cortex has sparse direct anatomical connectivity with the amygdala (79). Together, this pattern of connectivity alterations suggests that additional polysynaptic pathways may also contribute to observed vlPFC and dlPFC modulation of the amygdala and their association with anxiety symptoms.

### Network associations with depressive symptoms

Depressive symptoms were also associated with greater inhibitory influence from the vlPFC to the vmPFC. Processing of self-related stimuli is associated with activity of the default mode network (80, 81), particularly the vmPFC (82), whose activity has been shown to track the perceived veracity of self-referential statements (82). Given this role, increased inhibition from the vlPFC may reflect a mechanism for modifying deeply held negative beliefs. Notably, vmPFC activity and connectivity are commonly disrupted in depression and have been shown to predict treatment response (83, 84). During task-based studies, these vmPFC alterations have been interpreted as reflecting the suppression of affective and self-referential content that conflicts with task engagement (84, 85). Although vmPFC connectivity also predicts treatment response at rest (86), whether it generalises to contexts like cognitive restructuring, where vmPFC is engaged rather than suppressed, remains an open question.

### Limitations

While participants were trained in cognitive restructuring techniques, including active rebuttal, reinterpretation, and perspective shifting, we did not record which strategy was deployed on each trial. This information could provide clearer insight into the observed associations, given that different strategies engage distinct prefrontal regions (19). The strategies may also have varied systematically across groups and symptom profiles. Moreover, the sample comprised predominantly young female participants. Although this demographic profile is consistent with the epidemiology of depressive and anxiety disorders, and age and sex were adjusted for in all neuroimaging analyses, replication in more age- and sex-balanced samples is needed to establish the generalisability of these findings. While we use the terms "bottom-up" and "top-down" to describe interactions between the prefrontal cortex and amygdala, these labels are best considered heuristic rather than literal descriptions of their relationship. Consistent with contemporary models, both regions provide complementary contextual information that shapes neural activity through dynamic, recurrent interactions rather than a strictly hierarchical regulatory process. Our connectivity findings are consistent with this framework, although the specific computational roles of these reciprocal interactions warrant further investigation.

## Conclusions

Our findings illustrated that cognitive restructuring reduced the endorsement of negative statements in clinical participants, yet the underlying neurobiology diverged by symptom profile. Depressive and anxiety symptoms demonstrated differential associations with dlPFC function, downstream prefrontal–subcortical circuitry, as well as heart rate during cognitive restructuring, suggesting processes central to CBT may engage partially distinct mechanisms across symptom presentations. Longitudinal studies are needed to determine whether these neural signatures prospectively predict treatment response and inform the targeting of cognitive interventions.

## Data availability

Individual participant and group level effective connectivity data for this study are available at: https://github.com/alecJamieson/CNBT_depression_anxiety/tree/main/Data.

## Code availability

MATLAB and R scripts used to generate the 2^nd^-level results and associated figures of this study are available at: https://github.com/alecJamieson/CNBT_depression_anxiety.

## Author Contribution Statement

AJJ: Conceptualization, Formal analysis, Methodology, Writing - original draft, Visualization. TS: Conceptualization, Funding acquisition, Software, Writing - review & editing. CGD: Writing - review & editing. SI: Investigation, Methodology, Writing - review & editing. JAA: Investigation, Writing - review & editing. BAM: Funding acquisition, Writing - review & editing. RKG: Investigation; Writing - review & editing. KLF: Resources, Writing - review & editing. BJH: Conceptualization, Funding acquisition, Methodology, Supervision.

## Acknowledgements

The authors thank Braden Thai, Holly Carey and Amy Nielson for their contributions to data collection. The authors also acknowledge the facilities and scientific and technical assistance of the National Imaging Facility, a National Collaborative Research Infrastructure Strategy (NCRIS) capability, at the Melbourne Brain Centre Imaging Unit, University of Melbourne. In addition, the authors are grateful to Siemens for providing the MP2RAGE sequence as a “works in progress package” and CMRR (University of Minnesota) for sharing the multiband EPI sequence.

