## Supplementary Materials for "Mapping the neural circuitry of cognitive restructuring in depressive and anxiety disorders"

**Supplementary Methods..………………………………………...…………………………2**

**Supplementary Results…………………………………………...………………………….5**

**Supplementary Tables……………………………………………………………………….8**

**Supplementary Figures…………………………………………………………………….16**

**Supplementary Methods**

**Image acquisition**

MRI acquisition was performed on a Siemens 7T Plus research scanner (Siemens Healthcare, Erlangen, Germany) equipped with a 32Rx/1Tx channel head coil (Nova Medical Inc., Wilmington MA, USA). Functional (T2*-weighted) images were obtained using a multi-band and GRAPPA accelerated gradient-echo planar imaging sequence (1) in the steady state (multi-band factor, 6; parallel acceleration factor, 2; repetition time, 800 ms; echo time, 22.2 ms; flip angle, 45°) in a 20.8 cm field-of-view, with a 130×130-pixel matrix, a slice thickness of 1.6 mm (no gap) and in-plane voxel size=1.6×1.6 mm. Eighty-four interleaved slices were acquired parallel to the anterior-posterior commissure line, covering the whole brain. The cognitive restructuring paradigm lasted 8 minutes and 22 seconds, corresponding to 628 whole-brain echo-planar imaging volumes.

A high resolution structural T1-weighted image was obtained from each participant using magnetization-prepared 2 rapid gradient echo sequence (MP2RAGE; 2) for co-registration with the functional images (parallel reduction factor, 4; repetition time, 5 seconds; echo time, 2.04 ms; flip angle, 13°) in a 24 cm field of view, with a 330×330–pixel matrix, in-plane voxel size = 0.75×0.75 mm, slice thickness = 0.75 mm with 224 sagittal slices aligned parallel to the midline. To minimise head movement during scanning, foam pads were inserted on either side of the participants’ heads. Respiration and cardiac pulse were also recorded during the session at 50 Hz and 200 Hz, respectively, using a respiratory belt and pulse-oximeter (Siemens, Germany) for subsequent physiological noise correction.

**Preprocessing**

Imaging data were preprocessed using Statistical Parametric Mapping (SPM) 12 (v7771, Wellcome Trust Centre for Neuroimaging, London, UK) within a MATLAB 2023b environment (The MathWorks Inc., Natick, MA) on the Spartan High Performance Computer hosted at The University of Melbourne (3). Motion artifacts were corrected by realigning each participant’s timeseries to the mean image, and all images were resampled using 4th Degree B-Spline interpolation. Participants were excluded if movement exceeded a mean total displacement of 1.6mm (~1 native voxel), as assessed through the Motion Fingerprint toolbox (4).

Each participant’s anatomical images (denoised MP2RAGE (5)) were co-registered to their respective mean functional image, segmented, and normalised to the International Consortium of Brain Mapping template using the unified segmentation and the Diffeomorphic Anatomical Registration Through Exponentiated Lie Algebra approach (6). To preserve spatial specificity, smoothing was applied to functional images with a 3.2 mm full-width-at-half-maximum Gaussian kernel. Physiological noise was modelled using the PhysIO toolbox (7). Cardiac and respiratory recordings were imported into the toolbox, in which physiological noise models were applied to these recordings. Normalised white matter and cerebrospinal fluid tissue segmentations from each participant's structural image were used to construct noise regions of interest from which mean timeseries and principal components were extracted as nuisance regressors using CompCor (8). A total of 24 physiological noise regressors were included as covariates in the first-level analysis. These included six cardiac, eight respiratory, four cardiac by respiratory interactions, one respiration response function, one cardiac response function, the top principal components and mean timeseries of the cerebrospinal fluid and white matter, and six motion regressors.

**Timeseries extraction, model estimation, and inference using parametric empirical Bayes**

To localise and identify peak coordinates of the cortical regions, we used the Challenge > Repeat contrast, whereas the Challenge < Repeat contrast was used to localise amygdala activity, consistent with its activation profile across these tasks. Given individual variability in amygdala lateralisation, a bilateral amygdala mask was used to extract its timeseries of all included voxels (9). The timeseries for our chosen regions of interest were then extracted at the single-subject level using the first eigenvariate of voxels within 4 mm of the subject-specific maxima that showed significant activation (p < 0.05). These subject-specific maxima were required to be within 8mm of the group level maxima. If a given region showed inadequate activation, the threshold was incrementally relaxed up to *P* < 0.5, consistent with current guidelines (10).

Model estimation followed an iterative procedure: each participant’s model was first inverted, then used to construct a second-level PEB model whose posterior estimates served as priors for re-estimation at the single-subject level. This iterative inversion avoids local optima and improves parameter estimation (11). Re-estimated subject-level parameter estimates were then taken to the second level to model between-group effects using PEB while accounting for individual estimated variance (12). Posterior probabilities were computed using the free-energy (with vs without) method, which compares model evidence when specific parameters are included versus excluded. Parameters with posterior probability > 0.95 were considered to have *strong* evidence (13). The PEB model included six regressors: average connectivity across participants, the effects of depressive and anxiety symptoms, as well as age, sex, and antidepressant medication use as covariates. All regressors were mean-centred. We then searched over nested PEB models using Bayesian model reduction, pruning parameters that did not contribute to the overall model evidence (14, 15). Bayesian model averaging was performed on these models after the final iteration to determine the strength of connections in the last Occam’s window of 256 models.

**Leave-one-out cross-validation**

In the context of the PEB framework (13), leave-one-out cross-validation (LOOCV) is used to determine whether the size of interindividual effects on parameters is sufficiently large to predict a variable of interest (i.e., a parameter’s predictive utility)(13). In this case, this was employed to determine whether identified parameters could predict depressive or anxiety symptom severity. To do so, a group-level PEB model was estimated for all participants while excluding one participant, then this PEB model was used to predict the left-out subject’s symptom severity. Predicted symptom severity was then correlated with the observed symptom severity. A significant correlation between the expected and observed values demonstrates that the effect size was sufficiently large to predict the left-out subjects' symptom severity above chance.

**Heart rate modelling**

Using the pulse oximetry data, heart rate was derived continuously across the task period and preprocessed using the PhysIO toolbox (7). Heart rate values were downsampled and extracted at the volume level and subsequently categorised into the Challenge and Repeat conditions. Each condition window included the 12-second period during which participants either repeated or challenged negative statements. Prior to analysis, quality assurance was performed. Subjects for whom more than 10% of heart rate values fell outside a physiologically plausible range (30–220 bpm) were excluded (2 participants). A total of 36096 datapoints were available for this analysis (16 datapoints [0–12 s at 0.8 s intervals, inclusive] × 2 conditions × 8 blocks × 141 participants). A linear mixed-effects model was conducted in R using *nlme (16, 17)* to estimate associations between task condition, individual differences in depressive and anxiety symptoms, and heart rate. Heart rate was specified as the dependent variable, while condition (Challenge vs Repeat) was included as a fixed effect. Subject identity was included as a random intercept and a random slope for time to account for between-subject variability in both baseline heart rate and the heart rate trajectory over time. To examine whether depressive and anxiety symptoms were associated with heart rate, both variables were included as continuous fixed effects. Depression and anxiety scores were z-scored before analysis to facilitate interpretation and comparability of effect sizes. To test whether symptom severity moderated task-related changes in heart rate, interaction terms between condition and each symptom variable (Condition × Depressive Symptoms, Condition × Anxiety Symptoms) were included in the model. The final model was thus specified as follows: HR ~ Condition + Depressive Symptoms + Anxiety Symptoms + Time (mean-centred) + Condition*Depression *+* Condition***Anxiety + (Time | Subject). We also ran a supplementary model including only clinical participants to test whether the Condition-by-symptom effects held in this subgroup.

**Supplementary Results**

**Behavioural results**

As highlighted in *Supplementary* *Fig. S3*, clinical participants endorsed most statements more strongly than healthy controls. Three statements did not differ between groups: “I commonly think that people want to take advantage of me”; “I usually believe that people want me to fail”; and “I frequently believe that people will hurt me in order to get what they need”. To examine associations with depression and anxiety symptoms, we correlated item endorsement levels with the DASS subscales (*Supplementary Fig. S7*). All symptom domains showed significant correlations with most items, except the same three statements, which exhibited inconsistent or small effects. Notably, the strongest associations across subscales were with depressive symptoms.

**Supplementary heart rate results**

To characterise these effects within the clinical group specifically, the same model was fit restricted to clinical participants only (Supplementary Table S5). There was a robust main effect of condition, such that heart rate was significantly lower during the Challenge condition relative to Repeat (*b* = −1.19, *t* = −15.97, *p* < .001). There was a significant main effect of time, indicating a gradual decrease in heart rate across the trial (*b* = −0.05, *t* = −3.36, *p* < .001). Neither depressive symptoms (*b* = 1.18, *t* = 0.64, *p* = .523) nor anxiety symptoms (*b* = 0.26, *t* = 0.15, *p* = .882) were significantly associated with heart rate. As in the primary analysis, both symptom dimensions significantly moderated the condition effect, but in opposing directions. Specifically, greater depressive symptoms were associated with a larger condition-related reduction in heart rate (*b* = −0.20, *t* = −3.07, *p* = .002), whereas greater anxiety symptoms were associated with an attenuated condition-related reduction (*b* = 0.20, *t* = 3.19, *p* = .001).

**Whole sample intrinsic connectivity results**

Average intrinsic connectivity across all participants revealed inhibitory influences from the preSMA to the dlPFC and vlPFC, as well as from the dlPFC to amygdala, vmPFC to dlPFC and vlPFC and amygdala to dlPFC (*Supplementary Fig. S5A*). Excitatory influences were observed from the dlPFC to preSMA, vlPFC, and vmPFC, from the vlPFC to preSMA, dlPFC, and amygdala, from the vmPFC to amygdala and from the amygdala to vmPFC.

Depressive symptoms were associated with reduced inhibition from the vmPFC to vlPFC (*Supplementary Fig. S5B* and *D*). Anxiety symptoms were associated with more widespread changes, including greater excitation from the vlPFC to amygdala and from the amygdala to vlPFC, and greater inhibition from the vmPFC to dlPFC. These symptoms were also associated with reduced self-inhibition of the preSMA and greater self-inhibition of the amygdala (*Supplementary Fig. S5C* and *E*).

**Clinical group connectivity results**

We additionally repeated our effective connectivity analyses within the clinical group alone. For intrinsic connectivity, the clinical group only analysis recapitulated the association between depressive symptoms and reduced inhibition from the vmPFC to vlPFC, while also identifying several additional associations, particularly involving connections originating from the preSMA and vlPFC (*Supplementary Fig. S6B*). Similarly, all anxiety related effects were replicated, with the exception of the weakest effects from the vlPFC to amygdala, while additional alterations implicated connections involving the amygdala and dlPFC (*Supplementary Fig. S6C*).

Consistent with the whole sample analysis, depressive symptoms were associated with reduced excitation from the preSMA to dlPFC during challenging (*Supplementary Fig. S6E*). However, rather than the greater inhibition from the vlPFC to vmPFC observed in the whole sample, the clinical group only analysis identified greater inhibition in the reverse direction, from the vmPFC to vlPFC. Similarly, anxiety symptoms were associated with greater inhibition from the vlPFC to amygdala, as observed in the whole sample, but also with reduced inhibition from the amygdala to vmPFC (*Supplementary Fig. S6F*).

**Leave-one-out cross-validation**

We next used LOOCV to test whether symptom-related connectivity effects were sufficiently robust to predict symptom severity in held-out participants. Across the full sample, anxiety symptom severity was predicted by intrinsic connectivity from the vlPFC to amygdala (*r* = 0.14, *p* = 0.045), as well as modulatory connectivity from the vlPFC to amygdala (*r* = 0.16, p = 0.025) and from the vmPFC to preSMA (*r* = 0.30, *p* < .001). None of the three parameters predicted depressive symptom severity.

Within the clinical group alone, anxiety symptom severity was predicted by intrinsic connectivity from the vlPFC to vmPFC (*r* = 0.38, *p* < .001) and modulatory connectivity from the vlPFC to amygdala (*r* = 0.21, *p* = 0.036). Intrinsic connectivity from the vlPFC to vmPFC (*r* = 0.35, *p* = 0.001), intrinsic connectivity from the vlPFC to preSMA (*r* = 0.20, *p* = 0.045), and vlPFC self-connectivity (*r* = 0.27, *p* = .010) were shown to predict depressive symptom severity.

Supplementary Table S1.

*Contrast between the Challenge Condition and the Repeat Condition across all Participants.*

| Brain region | BA | Coordinates | | | Cluster size (1.6mm^3^ voxels) | *t* value |
| --- | --- | --- | --- | --- | --- | --- |
|  |  | X | Y | Z |  |  |
| Presupplementary motor area | 6 | −3 | 14 | 66 | 27068 | 16.71 |
|  | 6 | 0 | 13 | 58 |  | 14.98 |
| Dorsal anterior cingulate cortex | 32 | 2 | 18 | 45 |  | 13.32 |
| Cerebellum | – | 32 | −59 | −27 | 16259 | 10.88 |
| Posterior cingulate cortex | 23 | 5 | −51 | 29 |  | 10.31 |
|  | 23 | 3 | −56 | 22 |  | 10.14 |
| Caudate | – | 18 | 11 | 14 | 1570 | 10.35 |
|  | – | 18 | 5 | 21 |  | 9.28 |
|  | – | 18 | 14 | 5 |  | 7.31 |
| Temporal pole | 38 | 53 | 19 | −22 | 391 | 9.05 |
|  | 38 | 58 | 5 | −14 |  | 5.96 |
| Parahippocampal cortex | 36 | −14 | −24 | −13 | 553 | 7.47 |
| Periaqueductal grey | – | 0 | −30 | −13 |  | 6.38 |
| Pons | – | 0 | −32 | −22 |  | 4.81 |
| Angular gyrus | 39 | −40 | −59 | 27 | 979 | 7.15 |
|  | 39 | −40 | −64 | 37 |  | 5.24 |
|  | 39 | −51 | −62 | 22 |  | 5.20 |
| Visual association cortex | 18 | −6 | −93 | 21 | 864 | 7.05 |
| Cuneus | 19 | −5 | −86 | 29 |  | 6.55 |
|  | 18 | 0 | −86 | 16 |  | 5.55 |
| Cerebellum | – | −29 | −54 | −29 | 594 | 6.91 |
|  | – | −40 | −53 | −32 |  | 6.32 |
|  | – | −37 | −61 | −27 |  | 5.88 |
| Superior temporal gyrus | 22 | 46 | −34 | 3 | 229 | 6.54 |
| Thalamus | – | 19 | −13 | 11 | 95 | 5.91 |
| Parahippocampal cortex | 36 | 24 | −42 | −11 | 112 | 5.85 |
| Cerebellum | – | −14 | −37 | −16 | 52 | 5.72 |
| Cerebellum | – | −16 | −42 | −27 | 23 | 5.60 |
| Anterior cingulate cortex | 24 | −3 | 10 | 19 | 18 | 5.13 |
| Middle temporal gyrus | 21 | 58 | −13 | −13 | 80 | 4.88 |
|  | 21 | 64 | −2 | −18 |  | 4.84 |
| Putamen | – | 19 | 0 | 2 | 37 | 4.74 |
| Cerebellum | – | −19 | −32 | −30 | 11 | 4.69 |
| Middle cingulate cortex | 24 | −8 | −13 | 32 | 14 | 4.61 |
| Middle occipital gyrus | 19 | 51 | −69 | 11 | 24 | 4.48 |
| Superior occipital gyrus | 19 | −13 | −90 | 37 | 12 | 4.34 |
| Fusiform gyrus | 37 | 34 | −45 | −8 | 10 | 4.31 |
| Middle occipital gyrus | 19 | 45 | −67 | 2 | 19 | 4.07 |

*Note.* BA, Brodmann area. Coordinates are reported in MNI space. Results thresholded at *P*FDR < .001 (voxel-level). Clusters of ≥10 voxels are shown.

Supplementary Table S2.

*Contrast between the Repeat Condition and the Challenge Condition across all Participants.*

| Brain region | BA | Coordinates | | | Cluster size (1.6mm^3^ voxels) | *t* value |
| --- | --- | --- | --- | --- | --- | --- |
|  |  | X | Y | Z |  |  |
| Supramarginal gyrus | 40 | 56 | −35 | 43 | 54420 | 13.87 |
|  | 40 | 56 | −45 | 40 |  | 12.62 |
| Supramarginal gyrus | 40 | −62 | −38 | 40 |  | 12.38 |
| Inferior temporal gyrus | 37 | −54 | −59 | −6 | 5425 | 9.86 |
| Fusiform gyrus | 37 | −45 | −61 | −11 |  | 8.28 |
| Lingual gyrus | 18 | −8 | −75 | −5 |  | 7.10 |
| Orbitofrontal cortex | 11 | −24 | 30 | −13 | 624 | 7.42 |
|  | 11 | −32 | 50 | −16 |  | 5.87 |
|  | 11 | −32 | 42 | −16 |  | 5.56 |
| Dorsolateral prefrontal cortex | 46 | −45 | 37 | 13 | 696 | 7.39 |
|  | 46 | −50 | 43 | 14 |  | 7.33 |
|  | 46 | −45 | 40 | 21 |  | 6.11 |
| Primary visual cortex | 17 | 16 | −85 | 0 | 918 | 7.30 |
| Visual association cortex | 18 | 21 | −91 | −3 |  | 5.95 |
|  | 18 | 27 | −86 | −11 |  | 5.33 |
| Dorsomedial prefrontal cortex | 8 | 8 | 34 | 43 | 461 | 6.82 |
|  | 8 | 11 | 27 | 50 |  | 6.31 |
|  | 8 | 11 | 37 | 34 |  | 4.85 |
| Dorsolateral prefrontal cortex | 9 | −32 | 30 | 34 | 274 | 6.22 |
|  | 9 | −37 | 40 | 38 |  | 4.88 |
| Fusiform gyrus | 20 | 42 | −16 | −26 | 19 | 5.78 |
| Inferior temporal gyrus | 20 | −58 | −30 | −26 | 54 | 5.67 |
| Temporal pole | 38 | −27 | 11 | −45 | 62 | 5.62 |
| Thalamus | – | −10 | −29 | 6 | 20 | 5.55 |
| Superior frontal gyrus | 6 | 24 | 14 | 61 | 196 | 5.46 |
|  | 6 | 19 | 14 | 53 |  | 5.39 |
| Inferior temporal gyrus | 20 | −56 | −14 | −32 | 41 | 5.42 |
| Cerebellum | – | −27 | −69 | −42 | 104 | 5.40 |
|  | – | −22 | −69 | −50 |  | 4.13 |
| Thalamus | – | 16 | −24 | 8 | 20 | 5.27 |
| Middle cingulate cortex | 24 | 2 | 6 | 35 | 51 | 5.23 |
|  | 24 | 3 | −2 | 34 |  | 4.71 |
| Ventromedial prefrontal cortex | 11 | −8 | 26 | −21 | 14 | 5.07 |
| Middle temporal gyrus | 21 | 53 | 2 | −30 | 33 | 5.04 |
| Middle cingulate cortex | 24 | 6 | −11 | 35 | 15 | 4.94 |
| Inferior temporal gyrus | 20 | 58 | −14 | −30 | 44 | 4.87 |
| Amygdala | – | −21 | −3 | −19 | 52 | 4.79 |
| Temporal pole | 38 | −27 | 6 | −27 |  | 4.29 |
| Superior frontal gyrus | 9 | 19 | 35 | 32 | 91 | 4.66 |
|  | 9 | 24 | 43 | 34 |  | 4.61 |
| Postcentral gyrus | 2 | 38 | −27 | 48 | 22 | 4.65 |
| Pons | – | 11 | −26 | −27 | 18 | 4.62 |
| Cerebellum | – | −16 | −75 | −30 | 17 | 4.60 |
| Orbitofrontal cortex | 11 | −18 | 50 | −19 | 14 | 4.57 |
| Fusiform gyrus | 20 | 42 | −27 | −18 | 11 | 4.56 |
| Dorsomedial prefrontal cortex | 9 | −8 | 42 | 27 | 40 | 4.50 |
| Dorsomedial prefrontal cortex | 8 | 3 | 43 | 54 | 20 | 4.44 |
| Anterior prefrontal cortex | 10 | −34 | 56 | −5 | 22 | 4.38 |
| Paracentral lobule | 4 | 19 | −27 | 77 | 16 | 4.34 |
| Paracentral lobule | 4 | −19 | −30 | 77 | 25 | 4.33 |
| Ventromedial prefrontal cortex | 10 | 6 | 67 | −14 | 19 | 4.32 |
| Pregenual anterior cingulate cortex | 32 | 13 | 38 | 2 | 11 | 4.15 |
| Entorhinal cortex | 28 | 24 | 2 | −32 | 10 | 4.07 |
| Superior parietal lobule | 5 | 21 | −40 | 64 | 10 | 3.86 |

*Note.* BA, Brodmann area. Coordinates are reported in MNI space. Results thresholded at *P*FDR < .001 (voxel-level). Clusters of ≥10 voxels are shown.

Supplementary Table S3.

*Linear Mixed-effects Model Examining the influence of Condition, Time, Symptom Severity (Depression and Anxiety), and their Interactions with Condition on Heart Rate across all Participants.*

| Effect | Estimate | SE | df | *t* value | *p* value |
| --- | --- | --- | --- | --- | --- |
| Intercept | 71.42 | 1.03 | 35951 | 69.56 | < .001 |
| Condition (Chal vs Rep) | -1.18 | 0.04 | 35951 | -31.48 | < .001 |
| Depressive Symptoms (z) | 0.54 | 1.36 | 138 | 0.40 | .694 |
| Anxiety Symptoms (z) | -0.33 | 1.37 | 138 | -0.24 | .808 |
| Time (seconds) | -0.07 | 0.01 | 35950 | -6.23 | < .001 |
| Condition × Depression (z) | -0.22 | 0.05 | 35950 | -4.38 | < .001 |
| Condition × Anxiety (z) | 0.18 | 0.05 | 35950 | 3.65 | < .001 |

Supplementary Table S4.

*List of Negative Self-cognition Statements in the Cognitive Restructuring Paradigm.*

| 1. | I sometimes feel incompetent in the things I do |
| --- | --- |
| 2. | I often feel like I don’t measure up to others |
| 3. | I feel insignificant from time to time |
| 4. | I usually think other people are more competent than I am |
| 5. | I often think I will fail even if I make an effort |
| 6. | I occasionally believe that nobody will ever be attracted to me |
| 7. | I usually believe that I’ll be rejected if people discover my flaws |
| 8. | I repeatedly think that I’m a failure |
| 9. | From time to time, I think I am boring and uninteresting |
| 10. | I sometimes believe I am not good enough to be loved |
| 11. | I commonly think that people want to take advantage of me |
| 12. | I usually believe that people want me to fail |
| 13. | I often think I am incapable of changing my life |
| 14. | At times, I believe that I can’t do anything right |
| 15. | I occasionally think I have little value as a person |
| 16. | I frequently believe that people will hurt me in order to get what they need |

Supplementary Table S5.

*Linear Mixed-effects Model Examining the influence of Condition, Time, Symptom Severity (Depression and Anxiety), and their Interactions with Condition on Heart Rate in the Clinical Group.*

| Effect | Estimate | SE | df | *t* value | *p* value |
| --- | --- | --- | --- | --- | --- |
| Intercept | 69.85 | 2.07 | 18356 | 33.61 | < .001 |
| Condition (Chal vs Rep) | -1.19 | 0.07 | 18356 | -15.97 | < .001 |
| Depressive Symptoms (z) | 1.18 | 1.84 | 69 | 0.64 | .523 |
| Anxiety Symptoms (z) | 0.26 | 1.74 | 69 | 0.15 | .882 |
| Time (seconds) | -0.05 | 0.01 | 18356 | -3.36 | < .001 |
| Condition × Depression (z) | -0.20 | 0.07 | 18356 | -3.07 | .002 |
| Condition × Anxiety (z) | 0.20 | 0.06 | 18356 | 3.19 | .001 |

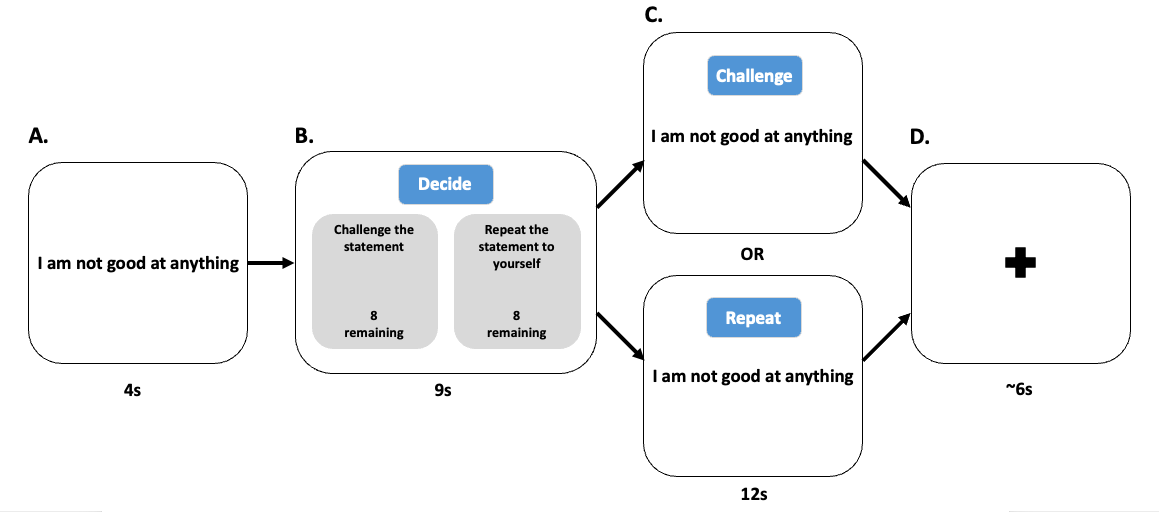

*Supplementary Fig. S1*. Schematic representation of the cognitive restructuring task. For each block, **A)** participants were first shown a negative statement for 4 seconds, then they had **B)** 9 seconds to decide whether to challenge the statement using Socratic questioning techniques or repeat the statement to themselves. After their choice, participants **C)** either cognitively restructured or repeated the statement for 12 seconds. Finally, **D)** a jittered fixation cross was shown for an average of 6 seconds.

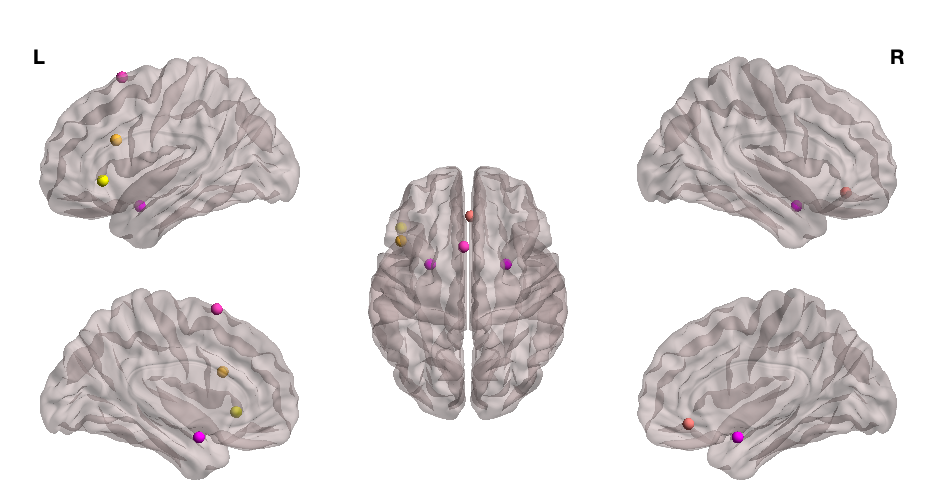

*Supplementary Fig. S2*. Visualisation of the brain regions used in the dynamic causal modelling analysis. To examine emotion regulation circuitry, we modelled directional interactions between the presupplementary motor area (pink), dorsolateral prefrontal cortex (dark yellow), ventrolateral prefrontal cortex (light yellow), ventromedial prefrontal cortex (light red), and amygdalae (purple). Render visualised using BrainNet Viewer (18).

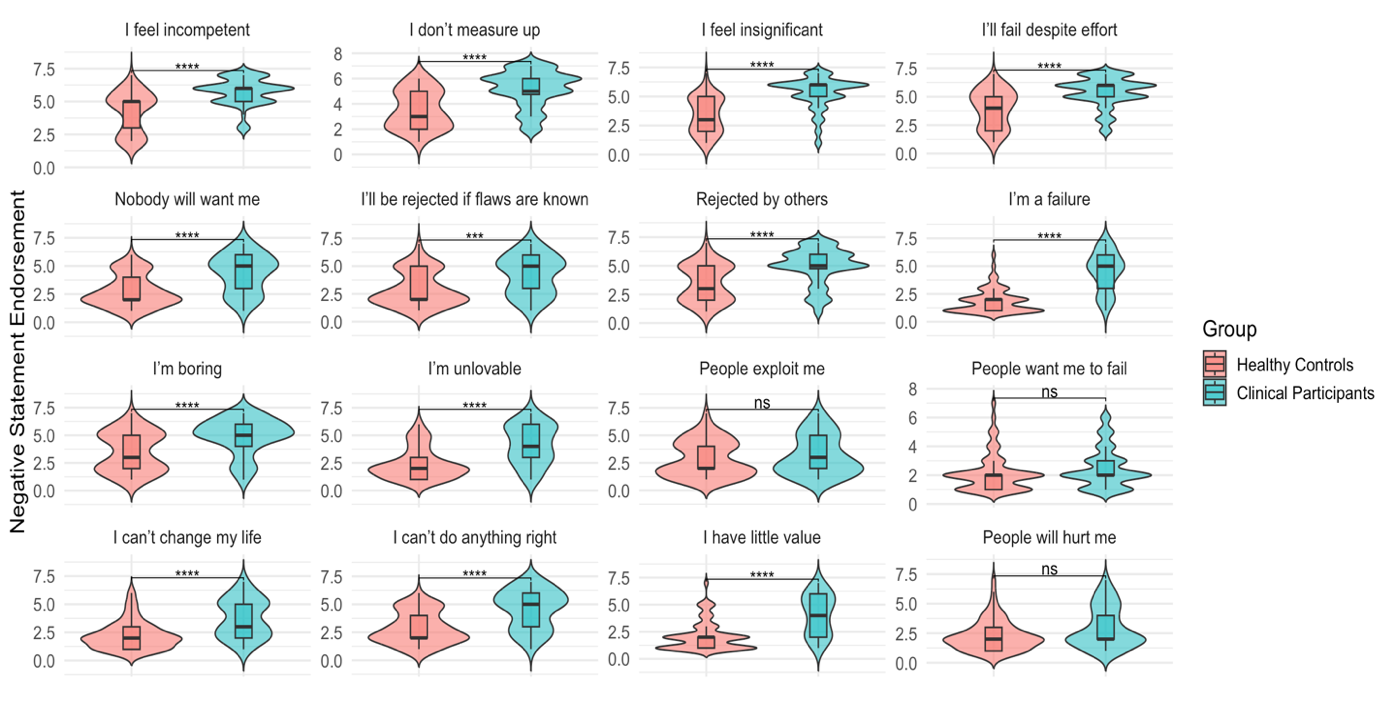

*Supplementary Fig. S3*. Violin plots depicting the distribution of endorsement levels across negative statements selected for the challenging condition of the restructuring task. Results are shown separately for healthy controls (salmon) and clinical participants (teal). Stars represent statistical significance after Bonferroni correction for multiple comparisons, with *** representing *p* < 0.001 and **** representing *p* < 0.0001. For the full list of the negative statements, see Supplementary Table S4.

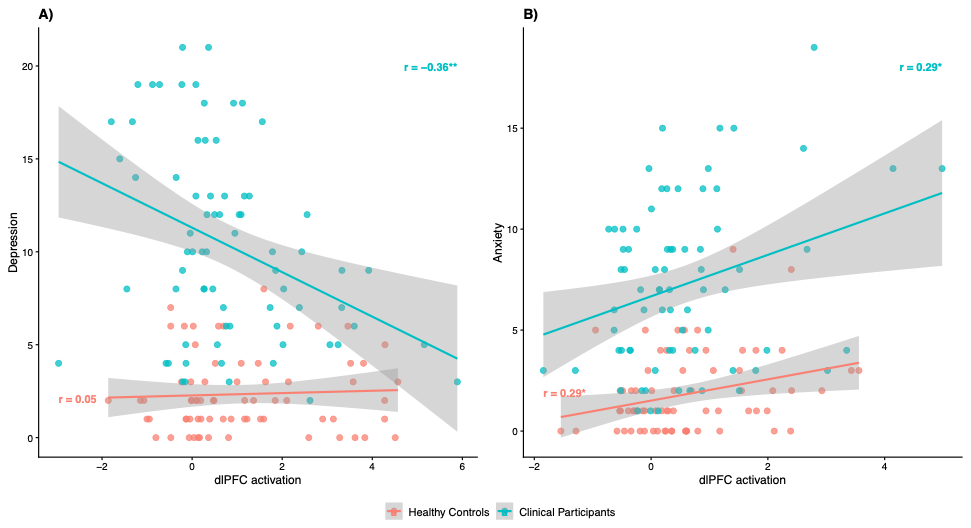
*Supplementary Fig. S4*. Associations between dlPFC activation and depression and anxiety symptoms in healthy and clinical participants. Regions of interest were extracted using the principal eigenvariate of the clusters implicated in the general linear modelling analyses investigating symptom severity (*Fig. 3*). Separate linear regressions were performed within healthy control and clinical participant groups to examine whether associations between dlPFC activation and symptom severity were driven by a specific group. **A)** Depression symptom effects were primarily driven by the clinical participant group, whereas **B)** anxiety symptom effects were consistent across healthy control and clinical participants. **p* < .05, ***p* < .01.

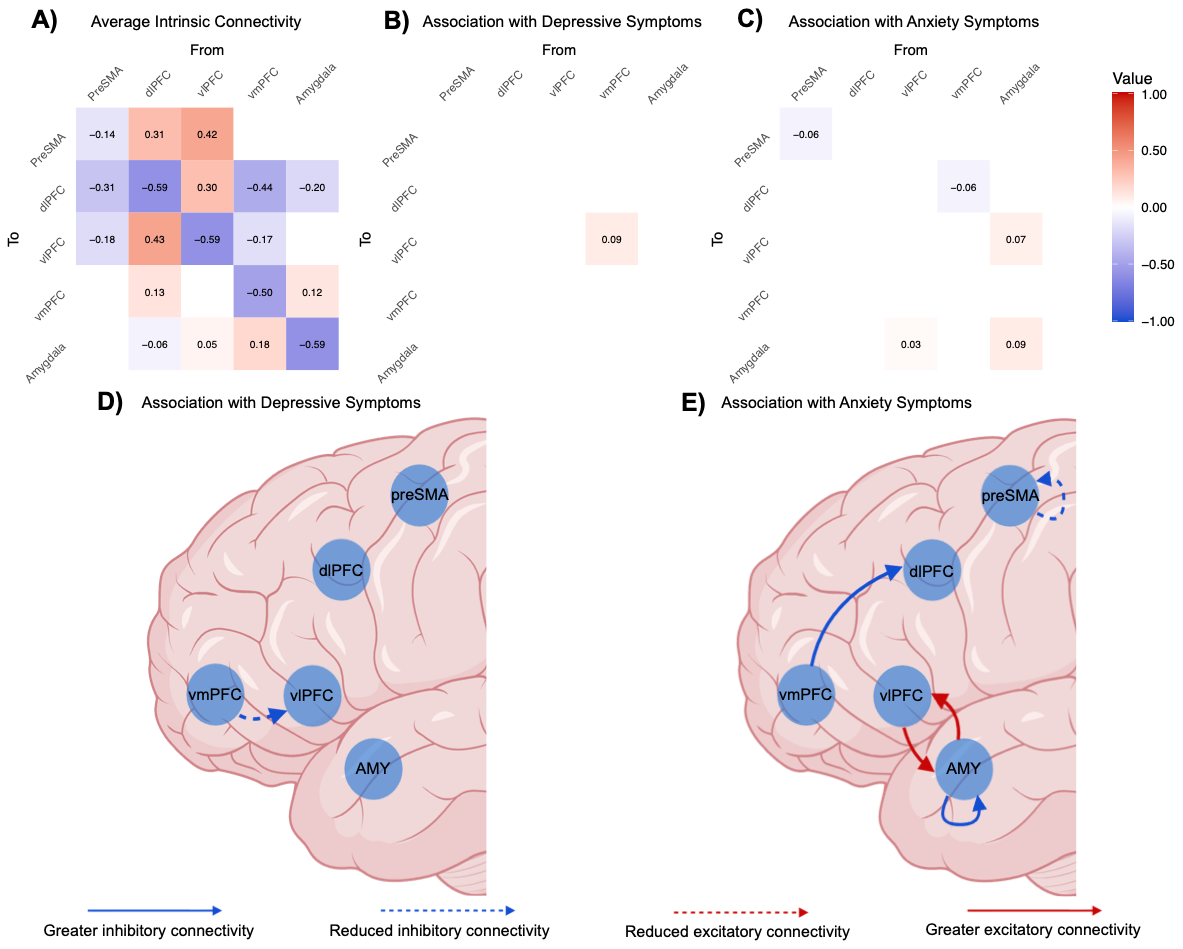

*Supplementary Fig. S5.* Intrinsic connectivity of emotion regulation regions across healthy controls and clinical participants. Adjacency matrices show the average effects observed in **A)** all participants, **B)** the effect of depressive symptoms, and **C)** the effect of anxiety symptoms. For **A)** red cells indicate excitatory connections whereas, blue cells indicate inhibitory connections. For **B)** and **C)** these values add (red) to or subtract (blue) from the connectivity in **A)**. Cells representing connectivity between two regions are measured in Hz and diagonal cells indicate inhibitory self-connections which are unitless log scaling parameters. **D)** and **E)** illustrate effective connectivity parameters associated with depressive and anxiety symptoms, respectively. Image created with BioRender ([www.biorender.com](http://www.biorender.com/)).

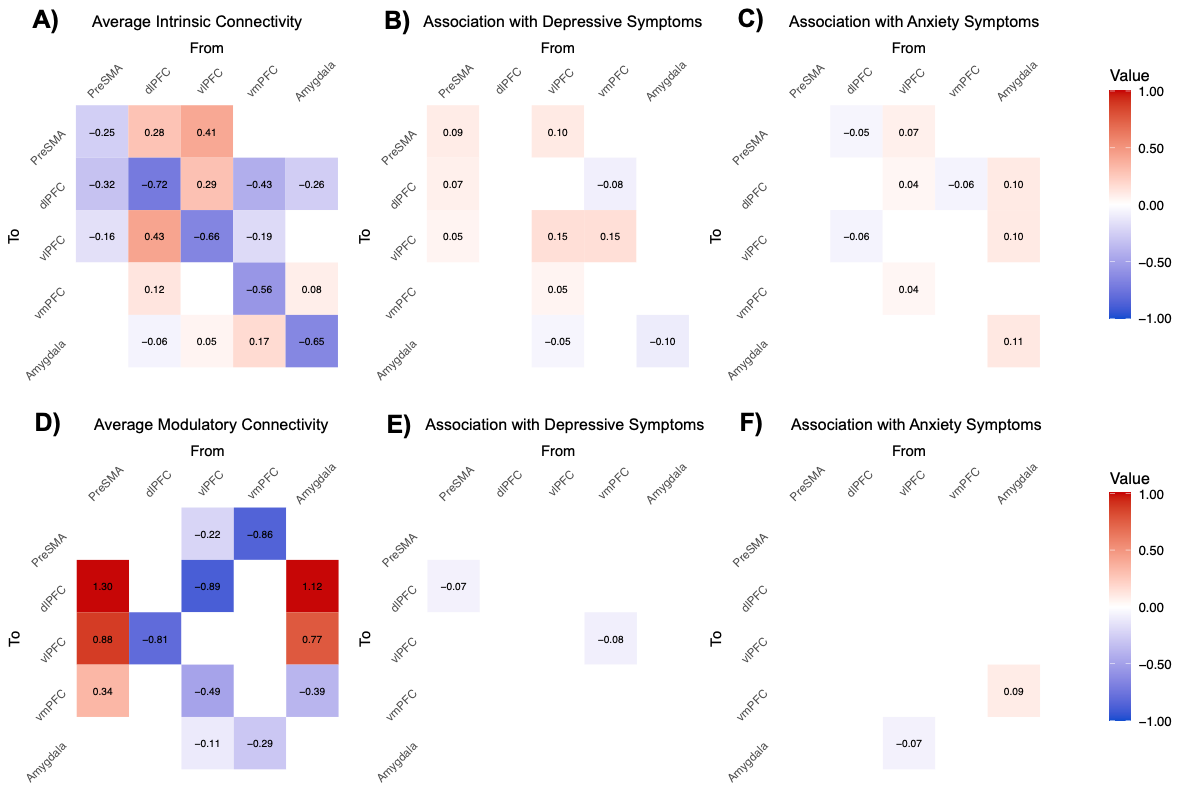

*Supplementary Fig. S6.* Intrinsic and modulatory connectivity of emotion regulation regions across clinical participants. Adjacency matrices show the average intrinsic effects in **A)** clinical participants, **B)** associated with depressive symptoms, and **C)** associated with anxiety symptoms. Corresponding matrices show the average modulatory effects in **D)** clinical participants, **E)** associated with depressive symptoms, and **F)** associated with anxiety symptoms. In **A)** and **D)**, red cells indicate excitatory connections and blue cells indicate inhibitory connections, whereas in **B)**, **C)**, **E)**, and **F)**, red and blue cells indicate increases and decreases, respectively, in connectivity relative to **A)** and **D)**. Off-diagonal cells represent connectivity between pairs of regions and are expressed in Hz, whereas diagonal cells represent inhibitory self-connections and are expressed as unitless log-scaling parameters.

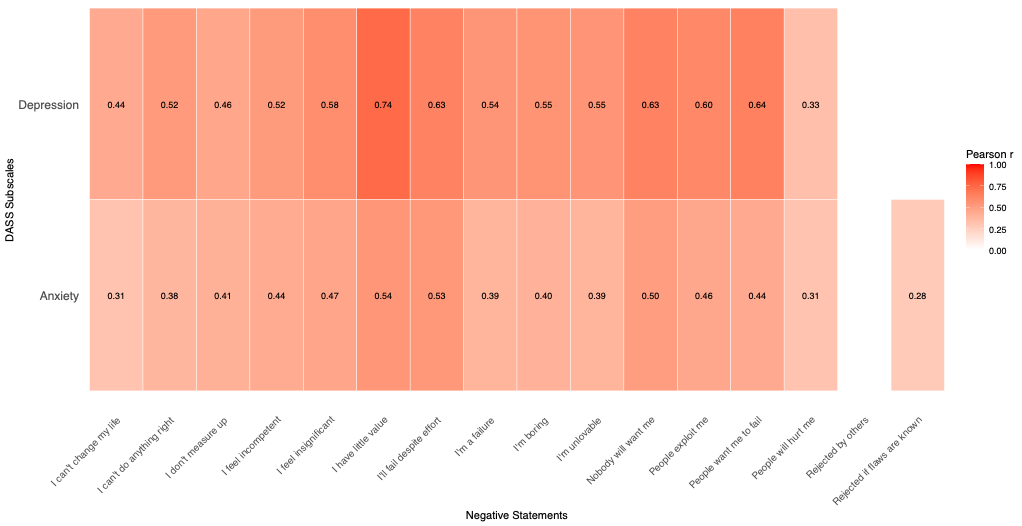
*Supplementary Fig. S7*. Pearson correlations of DASS depression and anxiety subscales with endorsement levels of individual negative statements. Results are Bonferroni-corrected.
